# Circadian clock regulates intestinal epithelial cell differentiation via NOTCH/Hes1 oscillations

**DOI:** 10.64898/2026.07.30.741736

**Authors:** Sevde Goker, Suengwon Lee, Bence Gaizer, Toru Matsu-Ura, Sabrina Tsoi, Gang Wu, Heewoong Lim, János Juhász, Attila Csikász-Nagy, Christian I Hong

## Abstract

The circadian clock regulates diverse cellular processes, including intestinal epithelial cell (IEC) proliferation. However, mechanisms regulating clock-dependent IEC differentiation remain unknown. We performed a time course RNA-Seq using the mouse small intestine and identified NOTCH signaling, a key mechanism regulating IEC differentiation, as one of the pathways under the control of circadian rhythms. Using mouse enteroids, we discovered that a NOTCH reporter, Hes1-luciferase, exhibits both ultradian or circadian oscillations depending on the stemness of mouse enteroids. Furthermore, single-cell analysis of Hes1-mCherry revealed that the period of *Hes1* oscillations varies widely, but circadian rhythms modulate the number of Hes1-mCherry+ cells in the population. Finally, we show that Paneth cell numbers fluctuate over the circadian cycle, suggesting that circadian clock-regulated *Hes1* drives the circadian dynamics of IEC composition. Our study provides a deeper insight into circadian regulation of IEC differentiation, which will be critical for applications of chronotherapies for digestive diseases.

## Introduction

The circadian clock is an endogenous timekeeping system that coordinates a variety of biological processes to align with the 24-hour day-night cycle. Light serves as the primary time cue, which is received by the retina and transmitted to the suprachiasmatic nucleus (SCN), master clock, located in the hypothalamus.^1,2^ The processed time information in the SCN is then sent to the peripheral oscillators and synchronizes them via neural and humoral pathways, thereby adapting the organism to its external environment.^3^ Although the SCN plays a central role in synchronizing the peripheral clocks, they can function independently of the SCN.^4,5,6,7^ In mammals, most cells possess their own molecular clock machinery.^8,9,10^ This molecular clock mechanism relies on a network of core clock genes and proteins that form a transcriptional-translational feedback loop (TTFL). The positive regulators, CLOCK and BMAL1, form a heterodimer promoting the expression of negative regulators, *Period* (*Per1, Per2, Per3*) and *Cryptochrome* (*Cry1, Cry2*), by binding the E-boxes on their promoters^11^. The protein products of *Per* and *Cry* genes then translocate back to the nucleus and inhibit the activity of CLOCK and BMAL1, which creates a time-delayed negative feedback loop generating autonomous oscillations with a period of approximately 24 hours.^12^ This molecular machinery also controls the expression of clock-controlled genes (CCGs) regulating numerous cellular, physiological and behavioral events, such as sleep-wake cycle, hormone release, and metabolism.^13^

Within the gastrointestinal system, the circadian clock plays a crucial role in maintaining homeostasis by regulating functions such as nutrient absorption, cell proliferation, and epithelial barrier integrity.^14,15^ The small intestine, the primary site for nutrient digestion and absorption, is lined by a single layer of intestinal epithelial cells (IECs) that undergo continuous renewal, with the entire lining being replaced every 4-5 days through a tightly regulated process of proliferation, differentiation, and cell shedding.^16^ Maintaining the balance between stem cell proliferation and differentiation is essential for proper intestinal function and homeostasis. A key player in this process is the NOTCH signaling pathway, which is important for the cell fate decision and the differentiation of IECs.^17,18,19^ Interestingly, emerging evidence suggests that the circadian clock, a fundamental mechanism that drives rhythmic physiological processes, is intimately involved in the regulation of intestinal homeostasis and epithelial cell turnover.^20,21,22,23^

NOTCH signaling pathway is essential for maintaining the balance between stem cell self-renewal and differentiation.^24^ NOTCH signaling uses lateral inhibition for cell-to-cell communication. While one cell (NOTCH-low cell) is providing the ligand (i.e., Delta), NOTCH receptor in its neighboring cell (NOTCH-high cell) binds to that ligand, which triggers a cascade that leads to the release of the NOTCH intracellular domain (NICD). NICD then translocates into the nucleus and binds to transcription factors such as RBPJ regulating transcription of target genes. Notably, NOTCH signaling activates the family of Hairy/Enhancer of Split (*Hes*) genes, which encodes transcriptional repressors that repress the expression of *Notch* and itself creating a negative feedback loop.^25^ This negative feedback loop enables oscillations of *Hes1*, which are observed in different contexts in mammalian tissues (e.g., developmental segmentation of vertebrae, neural progenitors).^26^

In the small intestine, NOTCH-activated HES1 represses another transcription factor, Atonal homolog 1 (ATOH1), which controls intestinal epithelial cell differentiation to secretory lineage.^27,28^ Consistent with this model, pharmacological inhibition of NOTCH signaling with gamma-secretase inhibitors or deletion of both *Notch1* and *Notch2* results in the conversion of other epithelial cell types to Goblet cells and increased number of Tuft cells.^29^ Furthermore, it was indicated that inactivation of HES1, HES3, and HES5 reduced cell proliferation and increased the number of secretory cells.^30^ In other others, high levels of *Hes1* expression promote the absorptive enterocyte lineage while suppressing secretory cell differentiation, and decreased *Hes1* expression allows the gene expressions for the differentiation of secretory cells, including goblet cells, enteroendocrine cells, and Paneth cells.^31^ By fine-tuning the levels of *Hes1*, NOTCH signaling ensures the proper composition and functionality of the intestinal epithelium. However, there have been no observations of circadian oscillations of *Hes1* oscillations to date.

Previously, it was demonstrated that the heterodimeric circadian clock transcription factor, CLOCK/BMAL1, binds to the E-box of *Notch2* and *Hes1* in epidermal stem cells.^32^ Despite the well-established roles of the circadian clock and NOTCH signaling in regulating various biological processes, the interaction between these two pathways in the context of intestinal epithelial cell differentiation remains largely unknown. In this study, we uncovered that NOTCH signaling pathway is one of the pathways that is controlled by circadian rhythms in the mouse small intestine. Then, using both bioluminescent and fluorescent NOTCH reporters, we discovered that circadian clock modulates the expression of *Hes1* in a population of IECs regulating their composition over the circadian cycle. By elucidating the molecular mechanisms underlying this interplay, this work deepens our understanding of the temporal regulation of intestinal epithelial homeostasis and its implications for gastrointestinal health.

## Results

### Approximately 29% of detectable transcripts are clock-controlled genes (CCGs) in the mouse small intestine

To obtain a global perspective of circadian regulation in the mouse small intestine, we performed a 48-hour time course with 2-hour resolution collecting jejunum for RNA-Seq (**Figure S1**). We used the RAIN algorithm,^33^ which identifies rhythmic patterns from time-series data, demonstrating 4,215 rhythmic genes out of 14,712 detectable transcripts (**Figure 1A**). This significant proportion of CCGs highlights the extensive regulation of the circadian clock on intestinal transcriptional activity, demonstrating approximately one-third of the transcriptome in the mouse small intestine is under clock control. Then, we performed phase set enrichment analysis (PSEA), which utilizes prior knowledge into the analysis of periodic data, to identify the signaling pathways that exhibit temporally coordinated expression profiles.^34^ PSEA revealed that numerous signaling pathways are under the control of circadian rhythms, and **figure 1B** shows representative clock-controlled signaling pathways across different circadian phases demonstrating that circadian rhythms orchestrate temporal organization of diverse signaling pathways ranging from cell cycle to immune response. Intriguingly, we observe ultradian and circadian rhythms of G1/S and G2/M scores, respectively (**Figure 1C**), indicating that the circadian clock controls G2/M transitions resulting in circadian time-dependent divisions in the small intestine. Our previous work demonstrated that, despite individual cell cycle times varying from 8 to 46 hours, the circadian clock in mouse enteroids orchestrate circadian time-dependent cell divisions *in vitro*.^35^ Based on this data, we speculate that the observed ultradian oscillations of G1/S score is due to the IECs with short cell cycle times in the intestine. Intriguingly, we discovered that genes involved in NOTCH signaling, which are known to regulate intestinal epithelial cell differentiation, are one of the clock-controlled signaling pathways (**Figure 1D**). However, molecular mechanisms regulating circadian clock-dependent IEC differentiation remain largely unknown.

**Figure 1.**
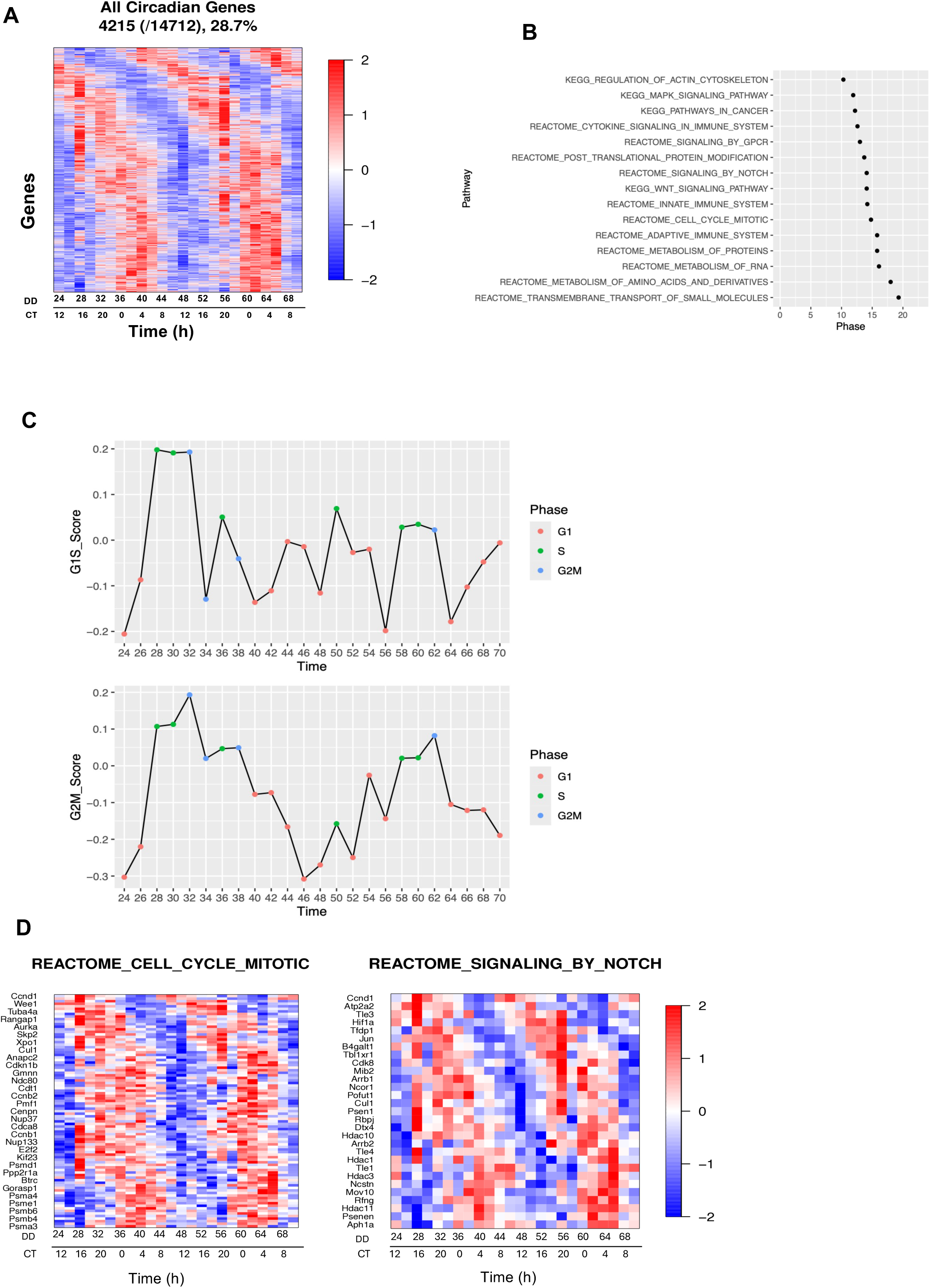
Approximately 29% of detectable transcripts are clock-controlled genes (CCGs) in the mouse small intestine. (A) Heatmap of circadian transcripts over 48 h in the mouse small intestine. (B) Representative circadian clock-controlled signaling pathways identified by phase set enrichment analysis (PSEA). (C) Cell cycle scoring analysis of G1/S score and G2/M score. (D) REACTOME Cell Cycle Mitotic and REACTOME Signaling by NOTCH Heatmaps demonstrating circadian-clock controlled genes involved in cell cycle and NOTCH signaling pathways. Data information: n=3 mice are used to generate RNA for each time point.

### Hes1-luc oscillations show ultradian or circadian rhythms depending on the stemness of mouse enteroids

The identification of NOTCH signaling as one of the clock-controlled signaling pathways, enables us to hypothesize that IEC circadian clocks regulate epithelial cell differentiation via NOTCH signaling. To test our hypothesis, we utilized mouse enteroids, 3-dimensional intestinal organoids derived from mouse crypts, to investigate molecular mechanisms and cellular dynamics in the small intestine, because it contains different cell types from stem cells to terminally differentiated IECs mimicking the crypt-villus architecture of the *in vivo* small intestine.^36,37^ And as a readout of NOTCH signaling, we employed Hes1-luc bioluminescent reporter system, because *Hes1* is one of the key target genes of NOTCH signaling.^30^

Growth and expansion of mouse enteroids involve regular changes of growth media every two days followed by passaging, which involves splitting of larger enteroids into smaller pieces, and subsequent seeding of dissociated enteroids in the Matrigel^TM^ with growth media. This physical splitting of enteroids promotes regeneration, expansion, and growth of enteroids. For circadian experiments, we administer 1-hour of dexamethasone (Dex) one day after passage, which resets the circadian clock of the population of enteroids and provides a reference phase of our circadian experiments. One day after passage, mouse enteroids show small, circular morphology without defined crypt and villus domains (**Figure 2A**). Under these conditions, we observe ultradian oscillations of Hes1-luc bioluminescent activity with an average period of 12.54 ± 5.16 hours (**Figure 2B**). Previously, we demonstrated disrupted circadian rhythms in both mouse and human intestinal enteroids under stem cell-enriched conditions,^35,38^ and indicated that the maturity of enteroids is critical for the development and function of enteroids,^39^ so we wondered whether maturation of mouse enteroids would alter ultradian oscillations of Hes1-luc reporter activity. To test this hypothesis, we allowed mouse enteroids to grow for three days after passage until we observed distinct morphology demonstrating both crypt and villus domains (**Figure 2C**). Intriguingly, Hes1-luc activity shows circadian oscillations with a period of 24.97 ± 1.79 hours when the bioluminescent imaging experiment was initiated 3 days after passage (**Figure 2D).**

**Figure 2:**
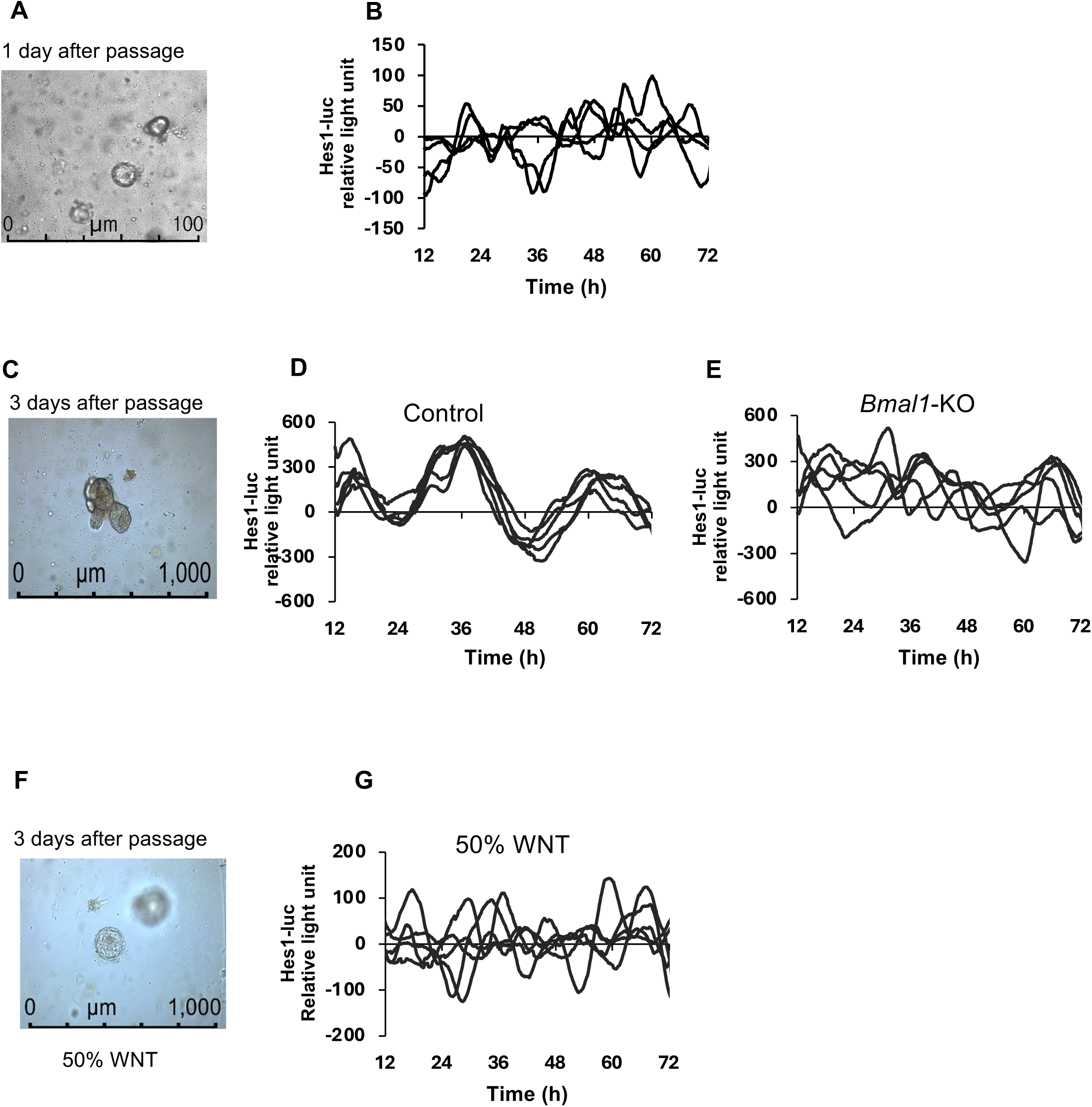
Hes1-luc oscillations exhibit either ultradian or circadian rhythms at the population level depending on the stemness of mouse enteroids. (A, C, and F) Representative images of mouse enteroids one day after passage in control conditions (A); three days after passage in control conditions (C); and three days after passage in stem cell-enriched (50% WNT) conditions (F). (B) Detrended bioluminescence data of Hes1-luc oscillations starting from one day after passage. Average period is 12.54 ± 5.16 hours. (D) Detrended bioluminescence data of Hes1-luc oscillations starting from three days after passage. Average period is 24.97 ± 1.79 hours. (E) Detrended bioluminescence data of Hes1-luc oscillations starting from three days after passage in *Bmal1*-KO enteroids. Average period is 19.89 ± 1.33 hours. (G) Detrended bioluminescence data of Hes1-luc oscillations starting from three days after passage under stem cell-enriched conditions. Average period is 14.6± 2.71 hours Data information: n≥4 technical replicates, X-axis indicates time post-Dex synchronization.

To test whether the IEC clocks regulate circadian rhythms of Hes1-luc activity, we used two independent conditions to disrupt circadian rhythms. First, we used *Bmal1*-KO enteroids derived from conditional *Bmal1*-KO mice (CagCRE;*Bmal1-f/f*) to genetically disrupt circadian rhythms, which showed a loss of circadian oscillations of Hes1-luc activity (**Figure 2E**). Second, we used 50% WNT-conditioned media to grow mouse enteroids in a stem cell-enriched condition **(Figure 2F)**, which results in dramatic reduction of amplitude of circadian rhythms.^35,38^ As expected, we observed that circadian oscillations of Hes1-luc activity are abolished under these conditions even though these experiments were performed three days after passage (**Figure 2E** **and** **Figure 2G**), indicating that the circadian rhythmicity of Hes1-luc expression relies on a functional circadian clock.

### Single-cell tracking reveals heterogeneous *Hes1* dynamics in mouse enteroids

Previous studies demonstrated ultradian rhythms of *Hes1* oscillations regulating numerous aspects of developmental processes ranging from vertebrate segmentation to embryonic neural stem cell differentiation.^25,40,41^ For example, the period of oscillations of HES1 and HES7 in the segmentation clock is approximately 2 and 4-6 hours in mouse and human embryos, respectively.^26,42^ However, we observe ultradian oscillations of Hes1-luc with a period of between 14.6 and 19.89 hours in the clock-disrupted enteroids, which is significantly longer than the previously observed period length. We speculated that this may be due to the fact that we have been assessing *Hes1* oscillation dynamics in a population measuring bioluminescence outputs from 50-100 mouse enteroids. Hence, we constructed Hes1-mCherry mouse enteroids (**Figure 3A, Video S1 and S2**) to determine the duration of Hes1-mCherry expression at single-cell level.

**Figure 3:**
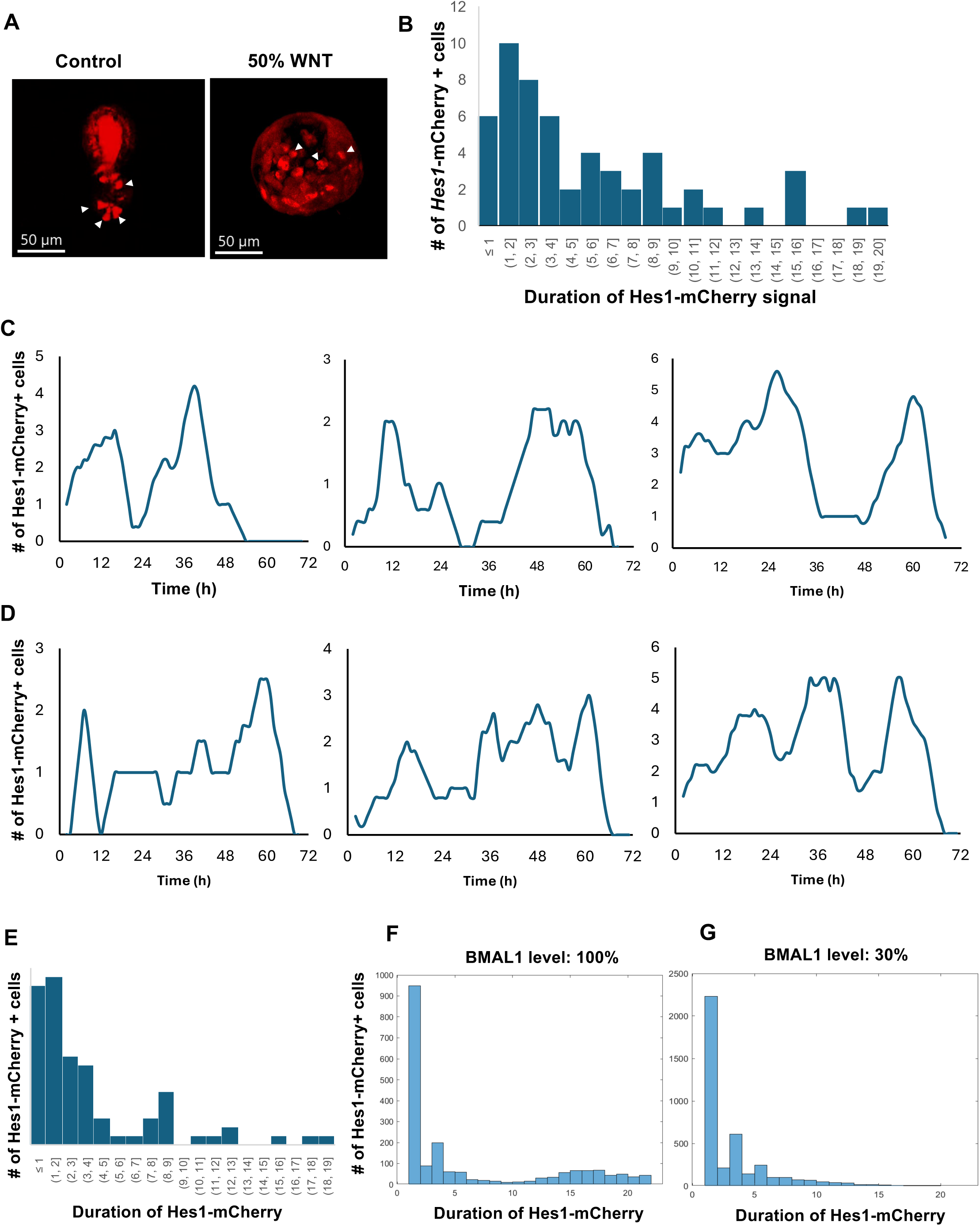
Single cell tracking reveals heterogeneous *Hes1* expression dynamics in mouse enteroids. (A) Representative image of Hes1-mCherry in control culture medium (left panel) or in stem cell-enriched 50% WNT-conditioned medium (right panel). White arrowheads, Hes1-mCherry+ cells. See also Supplementary videos 1 and 2. (B and E) Histogram of distribution of Hes1-mCherry signal duration in mouse enteroids cultured in control culture medium (B) or in 50% WNT-conditioned medium (E). n=55, n=77 Hes1-mCherry+ cells were analyzed in control vs. stem cell-enriched conditions, respectively, from three independent time courses. (C and D) Moving average graphs presenting number of Hes1-mCherry+ cells over time in control culture medium (C) or in 50% WNT-conditioned medium (D) from three independent time course experiments. (F and G) Histograms of Hes1-mCherry durations from stochastic simulations under 100% BMAL1 influence (control) (F) or 30% BMAL1 influence (disrupted clock conditions). Data information: n=3 technical replicates for each group, each experiment lasts over 60 hours.

Identical to the aforementioned experiments using Hes1-luc enteroids, we expanded Hes1-mCherry enteroids, administered 1-hour treatment of Dex to synchronize circadian rhythms, and performed time course confocal microscopy experiments starting from three days post-passage. Interestingly, we observe a wide range of Hes1-mCherry durations ranging from 1 to 20 hours (**Figure 3B**). However, a major portion of Hes1-mCherry+ cells (43.6%) show <3 hours of duration with a median period of 3.66 hours, which is consistent with the previously observed range of *Hes1* oscillations in the mouse segmentation clock and neural crest cells. And most recently, Waterings and colleagues showed a range of HES1 oscillation periods between 75 and 200 minutes of Hes1-Achilles reporter in mouse enteroids,^43^ which is also consistent with our data. However, the average duration of Hes1-mCherry expression is 5.46 hours, which is significantly longer than its median period, due to a significant number of Hes1-mCherry+ cells that show longer than 5 hours, which is 41.8% of the total number of analyzed cells. Then, how do these ultradian oscillations of *Hes1* in single cells manifest as circadian oscillations as a population?

To determine the potential circadian oscillations of Hes1-mCherry as a population, we measured the number of Hes1-mCherry+ cells over time. The moving average of Hes1-mCherry+ cells reveals that their overall population trend follows a longer cycle with an average period of 29.12 ± 5.26 hours (**Figure 3C**). Hes1-mCherry data does not show robust 24-hour oscillations due to the low number of Hes1-mCherry+ cells, where small variations/noises could alter the peak of oscillations. In contrast, we observe ultradian oscillations in the moving average of Hes1-mCherry+ cells with an average period of 16.7 ± 1.12 hours as a population in stem cell-enriched conditions (**Figure 3D**), which is consistent with our Hes1-luc bioluminescent activity data (**Figure 2E**). In stem cell-enriched conditions, we observe an increase of short-period (<3 hours) Hes1-mCherry+ cells (61%) with a median and average period of 2.48 and 3.91 hours, respectively (**Figure 3E**). This data is consistent with results from Weterings et al., where they show that shorter periods of Hes1-Achilles promote stemness and proliferative state.^43^ Furthermore, we observe a reduction in the number of Hes1-mCherry+ cells that show longer than 5 hours (23.4%) compared to the control (41.8%). We speculated that this difference may be due to the influence of circadian rhythms.

To obtain a deeper understanding of the *Hes1* oscillation dynamics in mouse enteroids, we constructed a mathematical model that links molecular mechanisms regulating *Hes1* oscillations^44^ and circadian clock machinery and performed stochastic simulations assessing *Hes1* dynamics (see supplementary information). For these simulations, we assumed that CLOCK/BMAL1 directly regulates the transcription of *Hes1* by binding to the E-boxes on the *Hes1* promoter, and the period of endogenous *Hes1* and circadian oscillations are 2 and 24 hours, respectively. When *Hes1* oscillator is coupled with circadian rhythms, our stochastic simulations qualitatively reproduce the observed histogram of *Hes1* dynamics with a majority of cells (53.15%) showing short duration (<3 hours) of *Hes1* and a significant portion (33.6%) of cells demonstrating >5 hours *Hes1* durations (**Figure 3F**). However, we observe a reduction of cells showing >5 hours Hes1-mCherry durations (19.36%) when we decrease the influence of BMAL1 on the *Hes1* transcriptions (**Figure 3G** and supplementary information) mimicking reduced amplitude circadian oscillations under stem cell-enriched conditions.^35,38^ Together, our data indicate that heterogeneous dynamics of *Hes1* duration is potentially due to the influence of circadian rhythms regulating the transcription of *Hes1*, and the circadian clock coordinates *Hes1* expression at the population level.

### Circadian clock regulates IEC composition in the mouse small intestine

Having established that the circadian clock modulates NOTCH signaling and *Hes1* expression dynamics, we next investigated whether this regulation extends to intestinal epithelial cell composition. Since NOTCH signaling plays a pivotal role in directing IEC fate,^31^ we hypothesized that circadian rhythms influence the distribution of differentiated cell types in the intestinal epithelium. To test this, we analyzed the expression of various IEC markers and assessed the rhythmicity of Paneth cell abundance over time. Our time-course RNA-seq data from the mouse small intestine shows that key IEC markers, *Muc13, Claudin4, Krt18, and Lyz1*, exhibit circadian oscillations **(Figure 4A).** Next, we wondered whether these transcriptional rhythms correspond to changes in specific epithelial cell populations, so as a proof of principle, we tested whether the number of Paneth cells changes over a circadian cycle. Paneth cells move downward after differentiation and provide nurturing and protection for the crypt base cells.^45^ They also express NOTCH ligands DLL1 and DLL4 which are necessary for NOTCH signaling activation,^46^ so the regulation of the differentiation into Paneth cells is critical for the small intestine homeostasis.

**Figure 4:**
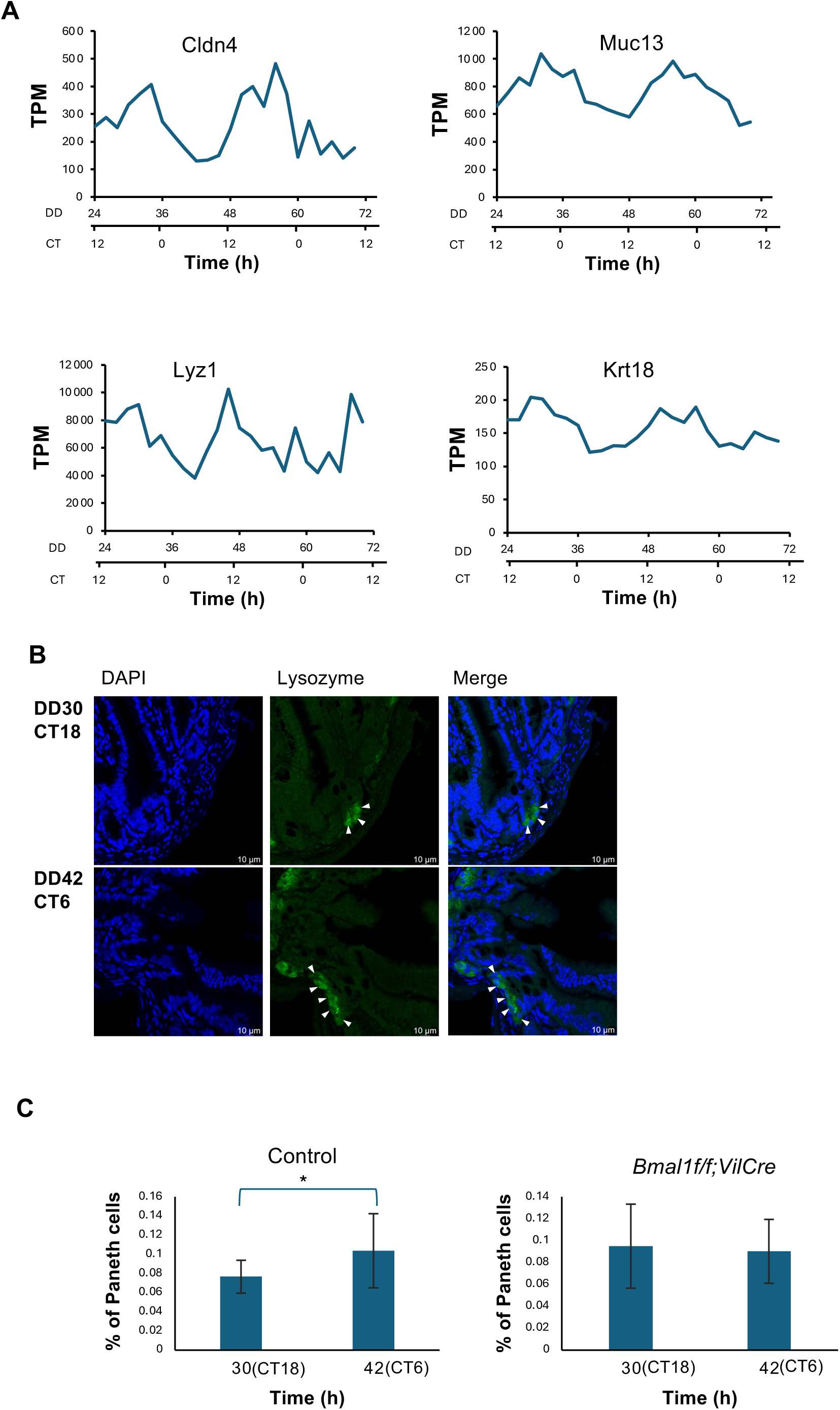
Circadian clock regulates IEC composition in the mouse small intestine. (A) Expressions of *Cldn4* (Enteroendocrine cells), *Muc13* (Goblet cells), *Krt18* (Tuft cells), and *Lyz1* (Paneth cells) over time from RNA-Seq data. TPM: transcripts per million; DD: Constant darkness; CT: Circadian Time (B) Representative images of DAPI (nuclear stain), Lysozyme (Paneth cell) and merge in mouse small intestine crypts. White arrowheads, Paneth cells. Scale bar, 50µm. (C) Average of Paneth cell abundance in different CTs in WT and *Bmal1f/f;VilCre* mouse small intestine crypts. Data information: Data represented as mean ± S.D. n≥3 technical replicates. 76 and 75 crypt samples in WT and *Bmal1f/f;VilCre,* respectively. *P<0.05 (two-tailed Student’s *t*-test)

To assess the number of Paneth cells, we performed immunostaining on mouse small intestinal tissue collected at two circadian time points (CTs) with a 12-hour interval **(Figure 4B).** Identical to our time course RNA-Seq experiment, C57B6 mice were entrained to 12:12 light:dark (LD) cycles for five days before transitioning to constant dark (DD) condition where their endogenous circadian rhythms are manifested in the absence of external time cues, and harvested mouse small intestine at DD30 (CT18) and DD42 (CT6). In the LD cycle, we monitor their diurnal rhythms based on Zeitgeber Times (ZT) where ZT0 and ZT12 indicate the beginning of light and dark phases. In DD, we keep track of CTs where CT0 and CT12 indicate the beginning of subjective day and subjective night, respectively. Interestingly, our quantification of Paneth cells revealed a higher percentage of Paneth cells at DD42 (CT6) compared to DD30 (CT18) **(Figure 4C).** In contrast, small intestine-specific *Bmal1-KO* mice (*Bmal1f/f;VilCre*) showed no significant difference in the number of Paneth cells between these two time points, indicating that the intestinal circadian clock is necessary for the observed time-dependent variation of Paneth cells. Together, these findings indicate that the circadian clock modulates IEC differentiation and epithelial composition by regulating NOTCH-dependent cell fate decisions over circadian cycles.

## Discussion

Our study provides novel insights into how the circadian clock regulates IEC differentiation through modulation of the NOTCH signaling pathway. By integrating transcriptomic, real-time bioluminescence recording, single-cell imaging approaches, and stochastic simulations, we demonstrate that *Hes1*, a key NOTCH target gene, exhibits dynamic rhythmic expression patterns in mouse enteroids, which are dependent on both the circadian clock and NOTCH signaling. Previous studies have established that pluripotent stem cells (PSCs) do not possess circadian rhythms, and the birth of circadian oscillation requires cellular differentiation from PSCs.^47,48^ In addition, we demonstrated that maturation of PSC-derived human intestinal organoids (HIOs) is required for robust oscillations of canonical clock genes regulating circadian clock-dependent functions.^39^ Importantly, we also demonstrated that the amplitude of circadian rhythms is dramatically reduced in stem cell-enriched conditions in both mouse and human intestinal enteroids suggesting a disruption of circadian rhythms in intestinal stem cells (ISCs) in mammals.^35,38^ In *Drosophila melanogaster*, however, circadian activity was recently reported in ISCs, which is downregulated during the differentiation process and re-emerged in totally differentiated cells.^49^ It was also previously shown that disruption of the circadian clock in enterocytes (ECs) dampens circadian clock activity in ISCs, which suggests that the differentiated cells positively regulate the circadian clock in ISCs in *Drosophila melanogaster.*^50^ In this report, we revealed that Hes1-luc expression shows ultradian and circadian profiles in stem cell-enriched and differentiated mouse enteroids as a population, respectively. These data suggest that there may be species-specific differences of circadian clock dynamics in intestinal stem cells.

The *Hes1* promoter contains several transcription factor binding motifs, which can be regulated by RBPJ, HES family proteins, and BMAL1. RBPJ regulates the expression of downstream target genes when it’s bound to NOTCH Intracellular Domain (NICD) recognizing (C/A/T)(G/A)TG(G/A/T)GAA motifs.^51^ In addition, HES proteins can form homo- and hetero-dimers with HES-related bHLH factors (e.g., HEY1) to repress *Hes1* expression by binding to the class C site (CACG(C/A)G) or the N box (CACNAG).^52,53^ And finally, *Hes1* promoter possesses 7 E-boxes (CANNTG), which can be regulated by a core circadian clock transcription factor, BMAL1.^32,54^. Initially, we hypothesized that BMAL1 could independently induce transcription of *Hes1*, but the *Hes1*(467RBPJ(-))-luc did not show any detectable bioluminescence (**Figure S2**) indicating that BMAL1 alone is not sufficient to induce the transcription of *Hes1*, and that Hes1-luc reporter indeed reflects NOTCH signaling-dependent activity via RBPJ binding motif. In the future, we plan to investigate complex transcriptional regulation of the *Hes1* promoter integrating circadian and NOTCH signaling-dependent transcription factors.

The period of *Hes1* oscillations is approximately 2 hours in the mouse segmentation clock,^44^ so we speculated that this period is too short for the intestinal circadian clock to phase-lock *Hes1* oscillations to 24 hours. Hence, we hypothesized that circadian rhythms might modulate *Hes1* expression at the population level. As predicted, our single-cell analysis of Hes1-mCherry signals in both control and stem cell-enriched conditions showed that the majority of Hes1-mCherry+ cells show short periods <3 hours (**Figure 3B**). Although individual cells exhibit short-period oscillations, the moving average of Hes1-mCherry+ cells revealed population-level trends of both circadian and ultradian rhythms under control and stem cell-enriched conditions, respectively, (**Figure 3**) indicating that the intestinal circadian clock modulates circadian expression of *Hes1* in a population of IECs. Furthermore, our stochastic simulations uncovered that the circadian clock is a likely mechanism underlying the emergence of the long-period (>5 hours) Hes1-mCherry+ cells (**Figure 3**). Interestingly, under stem cell-enriched conditions, the moving average of Hes1-mCherry+ cells exhibited a markedly longer ultradian rhythms (16.7 h ±1.12) at the population level, compared to that of individual cells (**Figure 3C and 3F**), suggesting the presence of an additional mechanism coordinating *Hes1* expression across the population.

Recently, Weterings and colleagues reported detailed oscillation dynamics of Hes1-Achilles in mouse enteroids demonstrating a range between 75 and 200 minutes,^43^ which is consistent with our data demonstrating <3 hours duration of Hes1-mCherry signals in the majority of cells (**Figure 3B**). Furthermore, they showed that lower periods of Hes1-Achilles are confined to near Paneth cells and lower-period oscillations promoted stemness, which is also consistent with our data demonstrating shorter duration of Hes1-mCherry in stem cell-enriched conditions (**Figure 3E**). Finally, they showed that the phases of Hes1-Achilles expression synchronized about 1.5 to 1 hour before mitosis. Intriguingly, our PSEA analysis of time course RNA-Seq data from the mouse small intestine revealed that the phase of NOTCH signaling occurs about 1 hour before the gene sets representing the mitotic cell cycle (**Figure 1B**). Together, these data suggest that the intestinal epithelial circadian clocks orchestrate the temporal organization of cell cycle and NOTCH signaling regulating circadian changes of IEC compositions including Paneth cells by temporally orchestrating lineage commitment and differentiation (**Figure 4**).

## Author contributions

S.G and C.I.H. designed the experiments. S.G., S.L., and T.M. performed the experiments. S.T. performed single cell analysis of Hes1-mCherry data. G.W. and H.L. performed bioinformatics analysis of RNA-Seq data. B.G., J.J., and A.C. performed and analyzed computer simulations. S.G., B.G., J.J., A.C., and C.I.H. wrote the manuscript, and C.I.H. supervised the overall project.

## Acknowledgements

We thank Dr. Ryoichiro Kageyama for pHes1(467)-luc and pHes1(467 RBPj(-))-luc plasmids, and Dr. Phillip Karpowicz for *Bmal1-f/f;VilCre mice*. We also thank Dr. Mari Kim for her assistance in time course experiment collecting the mouse small intestine. This work was supported by NIH R01 DK117005, R01GM156796 and University of Cincinnati Shaw’s Cancer Center Pilot Funding (C.I.H.). A.C. is supported by Hungarian National Research, Development and Innovation Office (NKFI/NRDI) through the Thematic Excellence Programe grant no. TKP2021-EGA-42. T.M. is supported by a grant from the Ministry of Education, Culture, Sports, Science and Technology of Japan to (19K08454 and 22K06071). We are grateful for imaging support from the University of Cincinnati Live Microscopy Core (NIH S10OD030402).

## Materials and Methods

| REAGENT or RESOURCE | SOURCE | IDENTIFIER |
| --- | --- | --- |
| Antibodies |  |  |
| Lysozyme rabbit polyclonal antibody for IF staining | Cell Marque | Cat 278A-17 |
| Goat Anti Rabbit secondary antibody | Thermo Fisher Scientific | Cat A-11034 |
| Chemicals, peptides, and recombinant proteins |  |  |
| PBS without Calcium and Magnesium | Corning | 21-040-CV |
| EDTA | Invitrogen | AM9260G |
| Advanced DMEM/F12 | Life Technologies | 12634-028 |
| Growth factor reduced Matrigel | Corning | CB-40230C |
| GlutaMAX | Gibco | 35050061 |
| HEPES | Gibco | 15630080 |
| Penicillin-Streptomycin | Gibco | 15140122 |
| N-2 Supplement (100X) | Gibco | 17502-048 |
| B-27 Supplement (50X) | Gibco | 17504-044 |
| Recombinant murine EGF | PeproTech | 315-09 |
| Nicotinamide | Sigma Aldrich | N0636 |
| Murine recombinant Wnt3a | R&D Systems | 1324-WN/CF |
| N-Acetyl-L-cysteine | Sigma Aldrich | A9165 |
| Gastrin I human | Sigma Aldrich | G9020 |
| A-83-01 | Sigma Aldrich | SML0788 |
| SB202 | Cayman Chemical | 10010399 |
| CHIR99021 | Cayman Chemical | 13122 |
| 0.25% Trypsin 2.21 nM EDTA | Corning | 25-053-CI |
| Polybrene | Millipore-Sigma | TR-1003-G |
| Y-27632 | R&D Systems | 1254/10 |
| Puromycin | Invivogen | ant-pr-1 |
| D-Luciferin (potassium salt) | Cayman Chemical | 14681 |
| Dexamethasone | Sigma Aldrich | D4902-500MG |
| DAPI | Thermo Fisher Scientific | 62247 |
| Citrisolve clearing reagent | Fisher Scientific | 22-143-975 |
| Trilogy | Cell Marque | CMX833 |
| 30% Hydrogen Peroxidase | ThermoFisher Scientific | 62247 |
| Critical commercial assays |  |  |
| RNeasy mini kit | Qiagen | 74106 |
| Experimental models: Organisms/strains |  |  |
| Mouse: C57BL/6 | The Jackson laboratory | 000664 |
| Mouse: Bmal1f/f | The Jackson laboratory | 007668 |
| Mouse: CAGCre-ER<sup>TM</sup> | The Jackson laboratory | 034795 |
| Mouse: Vil1-Cre | The Jackson laboratory | 021504 |
| Recombinant DNA |  |  |
| pRRLSIN_Hes1-467-Luc_SV40_puro (purchased from Addgene and modified as lentiviral plasmid) | Addgene | #41723 |
| pRRLSIN_Hes1-467-Luc_dRBPJ_SV40_puro (purchased from Addgene and modified as lentiviral plasmid) | Addgene | #43805 |
| pRRLSIN_Hes1-2500-mCherry_SV40_puro (purchased from Addgene and modified as lentiviral plasmid and luc tag replaced with mCherry tag) | Addgene | #43806 |
| Software and algorithms |  |  |
| R software | R 4.4.3 |  |
| Leica Application Suite X |  |  |
| Imaris Software | Imaris 10.2.0 |  |
| Other |  |  |
| BD 1-mL conventional insulin syringe | Fisher Scientific | 14-820-28 |
| 70 um cell strainer | Fisherbrand | 22363548 |
| Kronos Dio Luminometer Machine | ATTO Corporation | AB-2550 |

### Mouse

All animals were handled in accordance with IACUC protocols #23-02-10-01 (PI: Christian Hong, University of Cincinnati). We purchased C57BL/6 mice (Jackson lab: #000664) for WT controls, and generated the tamoxifen-inducible Bmal1 KO mouse (*Bmal1-f/f;CAGCreER*) and the small intestine-specific Bmal1-KO mice (*Bmal1-f/f;Vil-Cre*) by the breeding of *Bmal1-f/f* (Jackson lab: #007668) with *CAGCre-ER*^TM^ (Jackson lab: #0034795), and with *Vil1-Cre* (Jackson lab: #021504), respectively. Male mice with the age 6 to 14 weeks were used in all experiment including enteroid generation and immunostaining and RNA sequencing.

### Sample collection and RNA sequencing

For RNA sequencing to create a gene expression profile in the mouse small intestine, C57BL/6 mice (n=3 per time point) were initially entrained to 12-hour light/ 12-hour dark cycle for five days to synchronize their internal clocks. After 5 days, they were kept in constant darkness throughout the remainder of the experiment. Right after euthanizing the mice in CO2 chambers at certain time points, small intestine jejunum samples were collected every two hours for 46 hours and preserved by snap freezing for further RNA isolation procedure (Figure S1). After completion of the time course sample collection, RNA isolation was performed by using a RNeasy mini-kit column (Qiagen, 74106).

Isolated total RNA samples were shipped on dry ice and subjected to high-throughput sequencing using paired-end reads, generating ∼50 million reads per sample by Novogene. Raw RNA-sequencing FASTQ files were directly processed to generate expression files using the Kallisto^55^ embedded algorithm inside the Altanalyze module.^56^ Kallisto pseudo-aligned each transcript with the Ensemble reference transcriptome (Ensembl version 72) and calculated TPM (transcripts per million) by Altanalyze. For PSEA, the reference gene sets in c2.cp.v5.0.symbols.gmt obtained from the Molecular Signatures Database (MSigDB).

### Mouse small intestine crypt isolation and mouse enteroid generation

Crypt isolation, enteroid generation, and enteroid maintenance were performed as described in the previous publications.^57,58^

We used C57BL/6 and *Bmal1-f/f;CagCre*ER mice for crypt isolation to generate control and inducible clock disrupted mouse enteroids. Mice are euthanized in a CO₂ chamber for 10 minutes, followed by cervical dislocation after confirming that the mice are dead. The abdominal area is sterilized with 70% ethanol, then skin is cut off and approximately 3–4 inches of the jejunum (middle portion of the small intestine) are extracted. The intestinal segment is placed in a petri dish, flushed with cold PBS using a syringe to remove debris, and cut into three segments. Each segment is opened lengthwise, and the lumen is gently scraped to remove mucus and fecal matter. The scraped tissue samples are transferred to a clean 50 mL conical tube containing cold PBS, gently inverted for washing, and this washing process is repeated with another cold PBS containing canonical tube.

The cleaned samples were then transferred into a new canonical tube containing cold PBS with 5 mM EDTA and incubated on a shaker at 4°C for 30 minutes. During incubation, the tube is 1-minute vigorously shaken by hand every 10 minutes. Post-incubation, tissues and froth (if any) are removed, and the suspension is centrifuged at 800 rpm for 5 minutes at 4°C. The pellet is resuspended in 15 mL DMEM, centrifuged at 500 rpm for 5 minutes, and the supernatant is removed. The pellet is resuspended again in 5 mL DMEM and filtered through a 70 µm mesh into a new 50 mL canonical tube. The filtrate is centrifuged at 500 rpm for 3 minutes, and the resulting pellet is composed of crypt structures and used for enteroid generation.

Approximately 500 crypts per well are mixed with Matrigel and plated as 50 µL drops per well in a 24-well plate. Plates are incubated at 37°C for 30 minutes to solidify the Matrigel before adding 500 µL of pre-warmed media to each well. Media is replaced every 2 days.

### Maintenance mouse enteroids

Enteroids are passaged every 6–7 days to maintain and propagate them. Enteroids from four wells are collected into a 1.5 mL microcentrifuge tube, and Matrigel is broken down by pipetting. Samples are washed with cold PBS via a benchtop mini centrifuge for 1 minute to remove excess Matrigel and debris. The pellet of enteroids is resuspended in 0.7–0.75 mL PBS and mechanically dissociated by syringing 8–10 times with a syringe having 26½-gauge needle. After centrifugation with a benchtop mini centrifuge one more time, the supernatant is carefully removed, and after cooling down on top of ice, the pellet is mixed with Matrigel. The mixture is plated into a 24-well plate, incubated at 37°C for 30 minutes, and supplemented with 500 µL fresh pre-warmed media per well.

### Preparation of mouse enteroid culture medium

Mouse enteroids are cultured in Advanced DMEM/F-12 (Life Technologies, 12634-028) supplemented with 2mM GlutaMAX^™^ (Gibco, 35050061), 10 mM HEPES (Gibco, 15630080), 100 U/ml penicillin/100 μg/ml streptomycin (Corning, 30002CI), 1X N2 (Gibco, 17502–048), 1X B27 supplements (Gibco, 17504–044), 50 ng/mL EGF (PeproTech, 315–09) and 10% R-spondin1, 10% Noggin conditioned media generated in our lab.^59^

### Preparation of high WNT condition media

In order to promote stemness in the enteroid culture, we maintained them 50% WNT conditioned media which is made up of Wnt3a (R&D, 1324,WN/CF) supplemented with 10 mM Nicotinamide (Sigma Aldrich, N0636), 1 mM N-Acetyl-L-cysteine (Sigma Aldrich, A9165), 10 nM Gastrin (Sigma Aldrich, G9020), 500 nM A-83-01 (Sigma Aldrich, SML0788), 50ng/ml EGF (PeproTech, 315–09), 10 uM SB202 (Cayman Chemical, 10010399), 1X B27 supplements (Gibco, 17504–044), 20% R-spondin1, 10% Noggin conditioned media^59^ and Advanced DMEM/F-12 (Life Technologies, 12634-028) including 100 U/ml penicillin/100 μg/ml streptomycin (Corning, 30002CI), 2mM GlutaMAX^™^ (Gibco, 35050061), 10 mM HEPES (Gibco, 15630080).

### Lentiviral Transduction of mouse enteroids

For real-time monitoring *Hes1* reporter activity we performed Hes1-luc and Hes1-mCherry lentiviral transductions by following previously published protocol.^60^ For this purpose, plasmids were packaged into lentiviral vectors by the Viral Vector Facility at Cincinnati Children’s Hospital Medical Center (CCHMC). Briefly, after passaging the enteroids, we cultured them in mouse enteroid culture medium supplemented with 10 uM CHIR99021 - GSK-3 inhibitor- (Cayman Chemical, 13122) and 10mM Nicotinamide (Sigma Aldrich, N0636) to promote stemness condition, after a couple of days when the enteroids getting spherical formation, we collected 6 wells of them in a microcentrifuge tube, and wash for a few times to remove the Matrigel as much as possible. Then, 0.25 % Trypsin 2.21 nM EDTA (Corning, 25-053-CI) is added to the pellet and after vortex, the tube is incubated at 37 C for 3 minutes to get single cells. After incubation, 500 uL DMEM including 10% FBS is added into the tube in order to stop the trypsin effect. After spinning down for a minute, removed the supernatant and washed the pellet with 500 uL DMEM including 10% FBS, again. Then, the pellet is resuspended in mouse enteroid culture media including 8ug/mL Polybrene (Milipore-Sigma, TR-1003-G), 10 uM Y-27632 -ROCK inhibitor- (R&D Systems, 1254/10), 10 uM CHIR99021 - GSK-3 inhibitor-, and 50uL virus. 250 uL of this mixture is added into a well of 48-well plate, which is centrifuged at 600g for 1 hour. After centrifuge, the plate is incubated for 3 more hours at 37 C. Then, the digested enteroids are collected in microcentrifuge tube, washed with cold PBS, and mix with Matrigel to be seeded in 24 well. After 30 minutes incubation at 37 C, 500uL per well mouse enteroid media supplemented with 10uM ROCK inhibitor, 10uM CHIR, and 10mM Nicotinamide is added. After enteroids recovering, they are cultured in mouse enteroid media supplemented with 2 µg/ml Puromycin (Invivogen, ant-pr-1) for selection of enteroids carrying the interested reporter construct. This selection process is maintained for 2 weeks before further experiments.

### Bioluminescence recording

Hes1-luc expressing enteroids were monitored for bioluminescence over 72 hours, either starting from 1-day or 3-day post passage. The experimental procedures were performed as described previously.^38^ Transduced enteroids were embedded as 30 μL Matrigel domes in 35 mm culture dishes and maintained in 2 mL of mouse enteroid culture medium. For circadian synchronization, enteroids were treated with 100 nM dexamethasone for 1 hour. Following synchronization, the medium was replaced with 3 mL of mouse enteroid culture medium supplemented with 200 μM D-luciferin (Cayman Chemical, 14681), immediately prior to placement in the KronosDIO luminometer (ATTO). Bioluminescence signals were continuously recorded, and data analysis was performed using Kronos software, which provides time-series processing features such as moving average smoothing and signal detrending to facilitate the quantification of oscillatory dynamics.

### Time-course live imaging

Enteroids with Hes1-mCherry construction were generated to track Hes1 signal duration within single cells. Hes1-mCherry expressing enteroids were seeded into 8-well chamber slides with 20uL Matrigel drop per well and cultured in 250 μL either mouse enteroid culture media or high WNT-conditioned media. On the day of experiment, 1-hour 100nM Dexamethasone (Sigma Aldrich, D4902-500MG) treatment was performed for clock synchronization. After clock synchronization, media was replenished with 400 μL pre-warmed culture media before starting the live imaging. Live imaging of enteroids was performed for continuous 60-65 hours with 30 minutes intervals via Leica Stellaris 8 Confocal Microscope by using 10X objective. Excitation/emission settings were 405nm and 410-485 nm for DAPI; and 587nm and 595-700nm for mCherry.

### Single-cell analysis of Hes1-mCherry expression

After completion of time-course live imaging, the duration of Hes1-mCherry signal expression was analyzed using Imaris software. Hes1-mCherry expressing single cells were manually tracked to determine the duration of the signal. Only cells with clearly defined onset and offset time points were included in the analysis. Signals exhibiting unusually prolonged fluctuating expression patterns were excluded to ensure consistency and accuracy in quantification.

### Preparation for *in vivo* immunostaining

To assess the time-dependent abundance of Paneth cells, we collected small intestine jejunum samples with 12-hour intervals. C57BL/6 (n=4) and *Bmal1f/f;VilCre* (n=3) mice were used for this analysis. As explained in the RNA sample collection section, all mice were entrained under in 12-hour light/ 12-hour dark cycle, and tissue samples were collected at time points DD30 and DD42, corresponding to CT18 and CT6, respectively. Collected samples were fixed in 10% formalin at 4 °C overnight. Following fixation, tissues were processed, embedded in paraffin, and sectioned onto slides by the Integrated Pathology Research Facility at Cincinnati Children’s Hospital Medical Center (CCHMC).

### *In vivo* Paneth cell staining and cell counting

Slides were deparaffinized using Citrisolve clearing reagent (Fisher Scientific, 22-143-975) and hydrated in isopropanol for 5 minutes x 3 times. Then, antigen retrieval was performed using 1X Trilogy solution (Cell Marque, CMX833) in a pressure cooker, followed by blocking of endogenous peroxidase activity using 3% hydrogen peroxidase (Fisher Bioreagents, BP263-500). Samples were then incubating with blocking buffer (Ref) and stained with a primary Lysozyme antibody (Cell Marque, 278A-17) at 4 °C overnight. The following day, slides were washed with PBS and incubated with a secondary antibody (Life Technologies, A11034) for 2 hours at room temperature. Nuclear staining was performed using 1 μg/mL DAPI (ThermoFisher Scientific, 62247) for 2–3 minutes.

Fluorescent imaging was carried out using a Leica Stellaris 8 Confocal Microscope with 40X objective, and the excitation/emission settings were 405nm and 410-485 nm for DAPI; and 499nm and 505-565nm for Alexa488 (secondary antibody). For cell counting analysis, Imaris software is used and the ratio of Paneth cells per crypt region is decided by calculating the number of Lysozyme positive cells over total number of cells within each crypt region.

### Quantification and statistical analysis

All experiments were conducted with a minimum of three biological replicates. Periodicity of Hes1-luc and Hes1-mCherry oscillations were calculated via fast Fourier transformation.^35^ Comparison of Paneth cell abundance between two different time points was performed using unpaired two-tailed Student’s *t-*test assuming equal variances. Data are presented as mean ± standard deviation (SD), and statistical significance was defined as *p* < 0.05.

## List of Supplemental Information

Figure S1, Experimental design for *in vivo* time course

Figure S2, Comparison of Hes1-luc bioluminescence activity

Description of a mathematical model for stochastic simulations

Figure S3, Wiring diagram of the model

Table S1, Reactions of the Gillespie algorithm and corresponding parameters

