## Supplemental data for "Circadian clock regulates intestinal epithelial cell differentiation via NOTCH/Hes1 oscillations"

### List of Supplemental Information

Figure S1, Experimental design for *in vivo* time course

Figure S2, Comparison of Hes1-luc bioluminescence activity

Description of a mathematical model for stochastic simulations

Figure S3, Wiring diagram of the model

Table S1, Reactions of the Gillespie algorithm and corresponding parameters

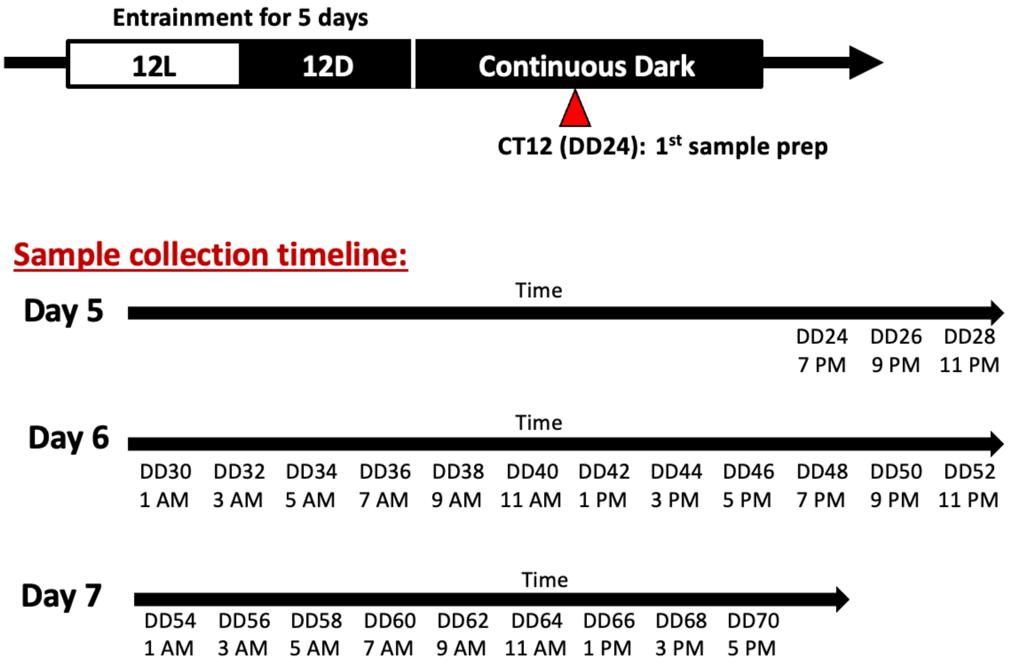


**Figure S1. Experimental design for *in vivo* time-course**

Entrainment lasted for 5 days under 12:12 light:dark cycles, then mice were kept under constant darkness throughout sample collection.

12L: 12 hours light; 12D: 12 hours dark; DD: constant darkness

DD24 corresponds to CT12 (starting time of subjective night)

n=3 mice for each time point

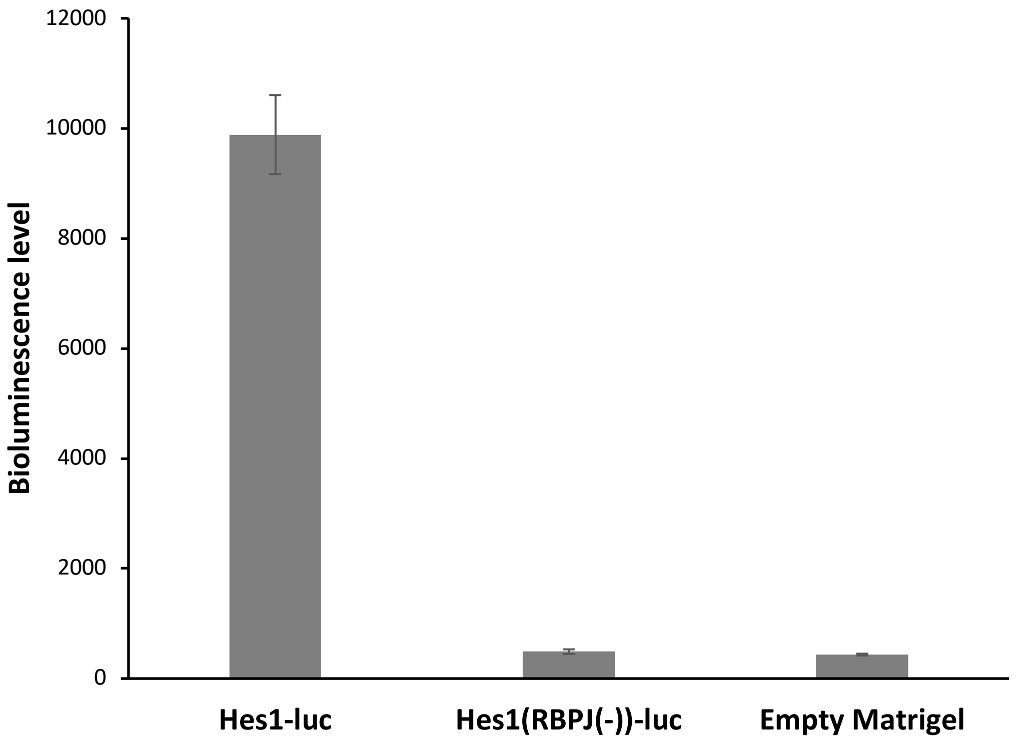


**Figure S2. Comparison of Hes1-luc bioluminescence activity**

Comparison of luminescence intensity level in mouse enteroids possessing Hes1-luc (control) or Hes1(RBPJ(-))-luc reporters and negative control (empty Matrigel).

Data information: Data represented as mean ± S.D. n≥3 technical replicates.

### Description of a mathematical model for stochastic simulations

Reconstructing oscillations of the *Hes1* autoregulatory network has been a widely researched topic over the last decades. For our simulations, we used a Hes1 model developed by Hirata and colleagues[1,2] and assume that BMAL1 directly binds to the E-boxes of the *Hes1* promoter activating transcription of *Hes1* (**Figure S3**).

Hirata et al. introduced of an additional molecule in their model, Interacting Factor (IF), which is downregulated by HES1 and forms complex with the Hes1 monomers. The addition of IF provides an additional complexity to the system that enables the system to efficiently reach stable limit cycles both in deterministic and stochastic simulations.

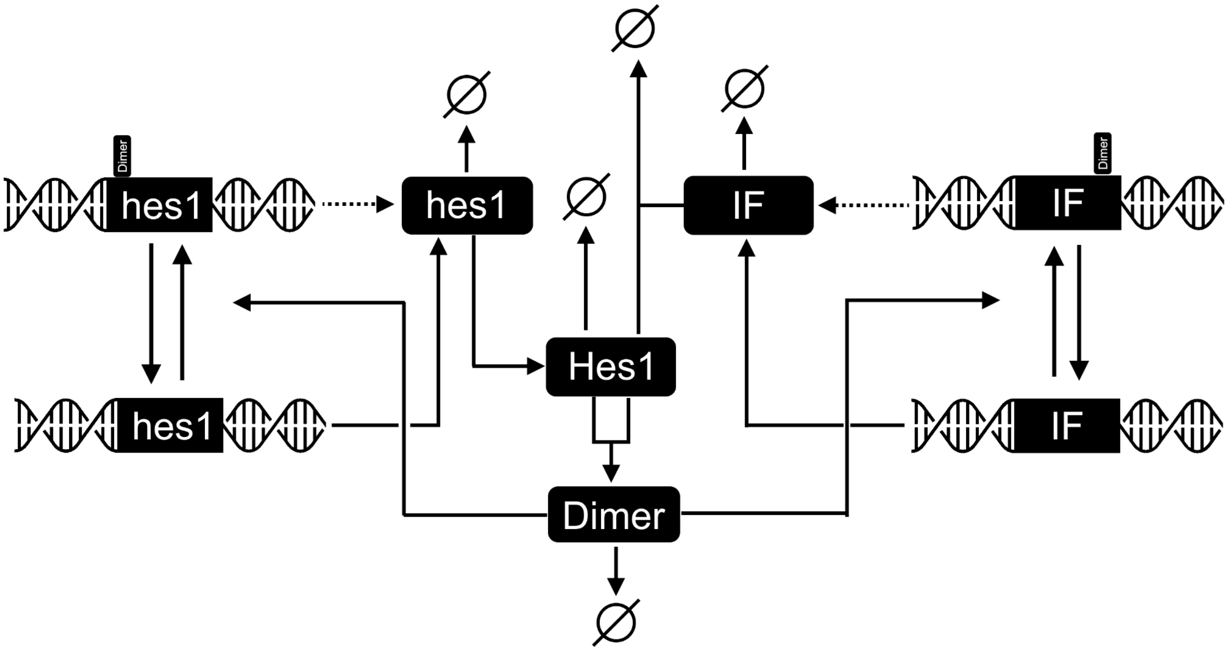


**Figure S3. Wiring diagram of the model**

First, used a deterministic model to find appropriate parameter sets and then constructed a stochastic model based on the Gillespie algorithm.[3] The model from Hirata et al. contains two Hill-type elements, both with a Hill-coefficient of 2. Since the possibility of employing Hill-type equations in a Gillespie stochastic model is debated, we followed the intuition that these equations generally model subsequent reactions and separated these equations; namely the occupation and liberation of promoters and the reduction in transcription of the respective genes. For the effect of the daily cycle on the Hes1 autoregulatory cycle we made some very simple assumptions. Interaction between the two systems is assumed to be one-way, therefore we use the level of BMAL1 as a fixed input on the system. We normalize its abundance and use a parameter to control its effect on the *Hes1* gene expression, modulating transcription of HES1-occupied and free promoters as well.

Following these considerations, our model consists of:

- Transcription of the *Hes1* mRNA
- BMAL1 directly activate transcription of Hes1 (Increase of H*es1* mRNA transcription by high BMALI1 activity)
- Translation of HES1 protein
- Synthesis of the IF
- Association between HES1 protein and IF (the created complex is assumed inactive.)
- Homodimerization of HES1
- Association of HES1 dimer to promoters of *Hes1* transcription and IF synthesis, reducing the activity of expression of both.

| Reaction No. | Reaction | Propensity | Parameters |
|---|---|---|---|

| | | | |
|---|---|---|---|
| 1 | $hes1 \rightarrow \emptyset$ | $\delta_h \cdot n_h$ | $\delta_h = 0.028\ \frac{1}{min}$ |
| 2 | $Hes1 \rightarrow \emptyset$ | $\delta_H \cdot n_H$ | $\delta_H = 0.108\ \frac{1}{min}$ |
| 3 | $Dimer \rightarrow \emptyset$ | $\delta_D \cdot n_D$ | $\delta_D = 0.036\ \frac{1}{min}$ |
| 4 | $IF \rightarrow \emptyset$ | $\delta_{IF} \cdot n_{IF}$ | $\delta_{IF} = 1.4\ \frac{1}{min}$ |
| 5 | $Hes1\ +\ IF \rightarrow \emptyset$ | $\frac{a}{K} \cdot n_H \cdot n_{IF}$ | $a = 0.03\ \frac{1}{nM \cdot min}$ |
| 6 | $hes1\ \rightarrow hes1\ +\ Hes1$ | $\beta \cdot n_h$ | $\beta = 1\ \frac{1}{min}$ |
| 7 | $Hes1 + Hes1 \rightarrow Dimer$ | $\frac{k_{dim}}{K} \cdot n_H \cdot (n_H - 1)$ | $k_{dim} = 1\ \frac{1}{nM \cdot min}$ |
| 8 | $hes1\ gene \rightarrow hes1\ gene\ +\ hes1$ | $\alpha_h \cdot n_{PH}$ | $\alpha_h = 3\ \frac{1}{min}$ |
| 9 | | $\frac{\alpha_h}{\gamma_h} \cdot (N_{PH} - n_{PH})$ | $\gamma_h = 100$ |
| 10 | $IF\ gene \rightarrow IF\ gene\ +\ IF$ | $\alpha_{IF} \cdot n_{PIF}$ | $\alpha_{IF} = 9000\ \frac{1}{min}$ |
| 11 | | $\frac{\alpha_{IF}}{\gamma_{IF}} \cdot (N_{PIF} - n_{PIF})$ | $\gamma_{IF} = 138$ |
| 12 | $free\ hes1\ promoter\ +\ Dimer \rightarrow occupied\ hes1\ promoter$ | $\frac{\lambda_h}{K} \cdot n_{PH} \cdot n_D$ | $\lambda_h = 0.25\ \frac{1}{nM \cdot min}$ |
| 13 | $occupied\ hes1\ promoter \rightarrow free\ hes1\ promoter$ | $\mu_h \cdot (N_{PH} - n_{PH})$ | $\mu_h = 0.096\ \frac{1}{min}$ |
| 14 | $free\ IF\ promoter\ +\ Dimer \rightarrow occupied\ IF\ promoter$ | $\frac{\lambda_{IF}}{K} \cdot n_{PIF} \cdot n_D$ | $\lambda_{IF} = 200\ \frac{1}{nM \cdot min}$ |

| | | | |
|---|---|---|---|
| 15 | $occupied\ IF\ promoter \rightarrow free\ IF\ promoter$ | $\mu_{IF} \cdot (N_{PIF} - n_{PIF})$ | $\mu_{IF} = 10\ \frac{1}{min}$ |

**Table S1 - Reactions of the Gillespie algorithm and corresponding parameters.**

System variables: $n_H$ - number of Hes1 protein molecules; $n_h$ - number of hes1 mRNA molecules; $n_{IF}$ - number of Interacting Factor molecules; $n_{PH}$ - number of free hes1 promoters; $n_{PIF}$ - number of free IF promoters; $n_D$ - number of Hes1 dimers. General parameters: $K = 6 * 10^{23} \cdot 10^{-12} \cdot 10^{-9} = 600$ - conversion ratio between molecule number and nanomolar; $N_{PH} = 30$ - total number of hes1 promoters; $N_{PIF} = 30$ - total number of IF promoters
